# Nucleocapsid protein of SARS-CoV-2 phase separates into RNA-rich polymerase-containing condensates

**DOI:** 10.1101/2020.06.18.160648

**Authors:** Adriana Savastano, Alain Ibáñez de Opakua, Marija Rankovic, Markus Zweckstetter

## Abstract

The etiologic agent of the Covid-19 pandemic is the severe acute respiratory syndrome coronavirus 2 (SARS-CoV-2). The viral membrane of SARS-CoV-2 surrounds a helical nucleocapsid in which the viral genome is encapsulated by the nucleocapsid protein. The nucleocapsid protein of SARS-CoV-2 is produced at high levels within infected cells, enhances the efficiency of viral RNA transcription and is essential for viral replication. Here we show that RNA induces cooperative liquid-liquid phase separation of the SARS-CoV-2 nucleocapsid protein. In agreement with its ability to phase separate in vitro, we show that the protein associates in cells with stress granules, cytoplasmic RNA/protein granules that form through liquid-liquid phase separation and are modulated by viruses to maximize replication efficiency. Liquid-liquid phase separation generates high-density protein/RNA condensates that recruit the RNA-dependent RNA polymerase complex of SARS-CoV-2 providing a mechanism for efficient transcription of viral RNA. Inhibition of RNA-induced phase separation of the nucleocapsid protein by small molecules or biologics thus can interfere with a key step in the SARS-CoV-2 replication cycle.

## INTRODUCTION

The etiologic agent of the Covid-19 pandemic is the severe acute respiratory syndrome coronavirus 2 (SARS-CoV-2) (https://www.who.int/emergencies/diseases/novel-coronavirus-2019). SARS-CoV-2 is an enveloped single-stranded, positive-sense RNA virus with a 30 kb genome, one of the largest among RNA viruses (Virology.org, 2020; Zhou et al., 2020). The viral membrane of SARS-CoV-2, which contains the spike protein, a glycoprotein and the envelope protein (Watanabe et al., 2020; Zhou et al., 2020), surrounds a helical nucleocapsid. In the nucleocapsid, the viral genome is encapsulated by the nucleocapsid protein and thereby protected from the host cell environment (McBride et al., 2014; Gralinski and Menachery, 2020). The nucleocapsid protein of human coronaviruses is produced at high levels within infected cells and is critical for virion assembly (McBride et al., 2014; Fehr and Perlman, 2015; Gralinski and Menachery, 2020; Liu et al., 2020). In addition, it enhances the efficiency of sub-genomic viral RNA transcription and is essential for viral replication (McBride et al., 2014). Because of its importance for diagnostic and therapeutic approaches to treat Covid-19 (Burbelo et al., 2020; Lu et al., 2020), there is an urgent need to define the molecular mechanisms that underlie the nucleocapsid protein’s fundamental viral role.

Liquid-liquid phase separation (LLPS) provides a highly cooperative mechanism to locally concentrate proteins and nucleic acids and promote cellular reactions (Forman-Kay et al., 2018; Alberti et al., 2019). Recent evidence indicates that negative-sense RNA viruses, which replicate in the cytoplasm of infected cells (Novoa et al., 2005), concentrate their replication machinery in dynamic compartments formed by LLPS of the viral structural proteins L, phosphoprotein (P) and nucleocapsid (N) protein (Nikolic et al., 2017; Heinrich et al., 2018). The genomes of positive-sense RNA viruses such as SARS-CoV-2, however, lack the genetic code for the phosphoprotein P, which is essential for LLPS in negative-sense RNA viruses (Nikolic et al., 2017; Heinrich et al., 2018; Guseva et al., 2020).

Here we investigate liquid-liquid phase separation of the nucleocapsid protein of SARS-CoV-2 and show that nucleocapsid protein LLPS concentrates components of the SARS-CoV-2 replication machinery providing a mechanism for enhanced viral transcription and replication.

## RESULTS

### LLPS of N^SARS-CoV-2^ and RNA into Protein/RNA-Dense Compartments

To investigate if the N protein of SARS-CoV-2 (further termed N^SARS-CoV-2^; Figure 1A) phase separates in the absence of other viral proteins, we measured the turbidity of N^SARS-CoV-2^ solutions at different protein concentrations. Up to 50 µM, the protein solution retained low absorbance (Figure 1B), despite its tendency to oligomerize (Ye et al., 2020). Next we tested LLPS of N^SARS-CoV-2^ in the presence of RNA. The 419-residue N^SARS-CoV-2^ contains an RNA-binding domain and a C-terminal dimerization domain embedded into long intrinsically disordered regions (Figure 1A). The globular domains as well as the intrinsically disordered regions of coronavirus N proteins bind to RNA (McBride et al., 2014). At 50 µM protein concentration, addition of 1 µM polyU (800 kDa) strongly increased turbidity (Figure 1B). Differential interference contrast (DIC) and fluorescent microscopy demonstrated the formation of spherical droplets (Figure 1C). The droplets contained both N^SARS-CoV-2^ and RNA (Figure 1C). N^SARS-CoV-2^/polyU droplets were robust against the presence of the aliphatic alcohol 1,6-hexanediol (Figure S1A). In contrast, addition of increasing amounts of NaCl dissolved the droplets (Figure S1B), indicating an important role of electrostatic interactions for RNA-induced LLPS of N^SARS-CoV-2^. Quantification of fluorescence intensities of N^SARS-CoV-2^ and RNA showed that their concentration is strongly increased inside droplets (Figure 1D), i.e. cooperative LLPS of N^SARS-CoV-2^ and RNA into protein/RNA-dense compartments occurs.

**Figure 1.**
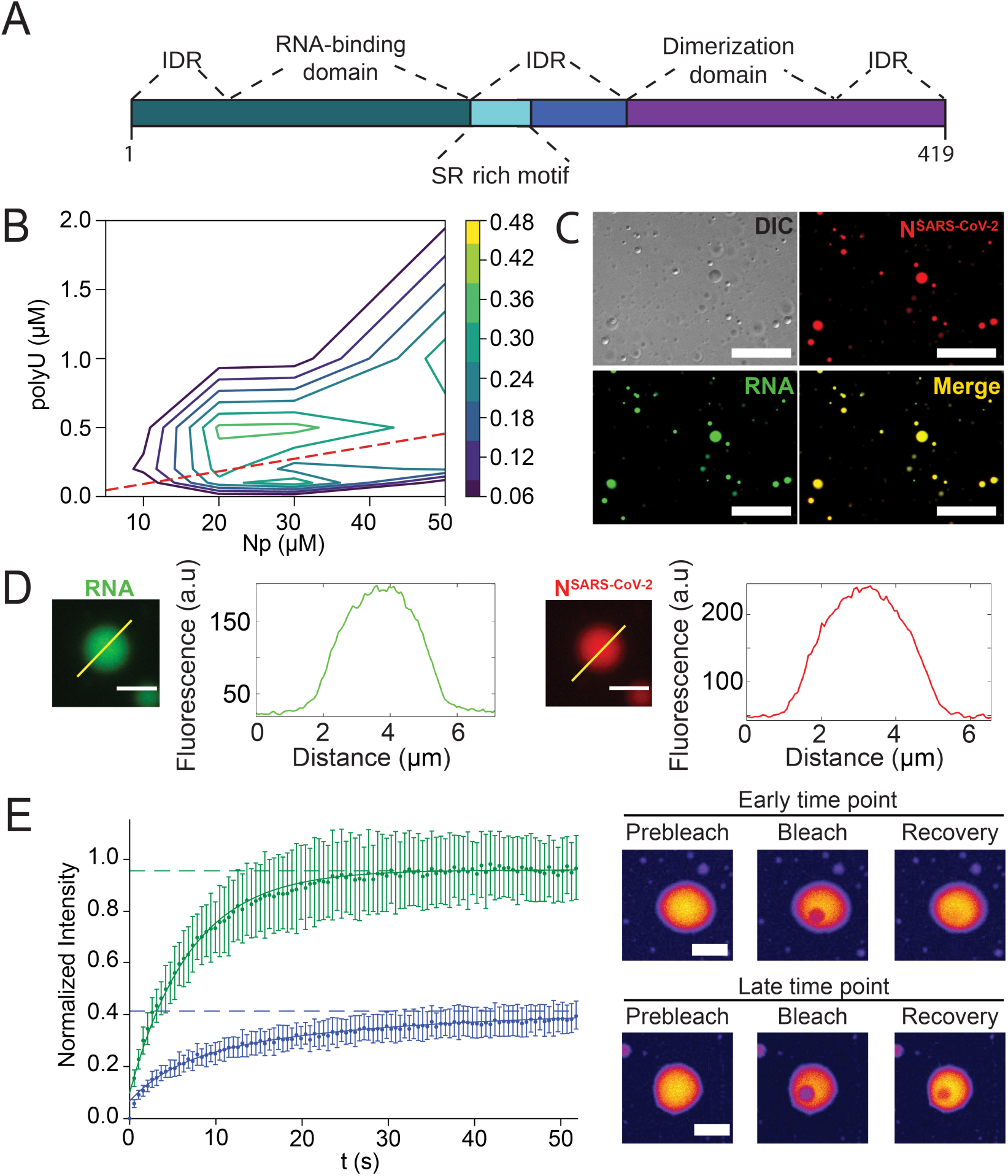
RNA-induced LLPS of the nucleocapsid protein of SARS-CoV-2. **(A)** Organization of N^SARS-CoV-2^ into two globular domains (RNA-binding domain and C-terminal dimerization domain) surrounded by long intrinsically disordered regions (IDR). The serine/arginine (SR)-rich region is conserved in coronaviruses. **(B)** Influence of RNA and protein concentration on N^SARS-CoV-2^/polyU-LLPS in 20 mM NaPi, pH 7.5, monitored by solution turbidity at 350 nm. Average values from three independent measurements are shown. The dashed line marks N^SARS-CoV-2^/polyU-concentrations at which charge neutralization occurs, assuming a charge of −1 per phosphate group. See also Figure S1. **(C)** Fluorescence and DIC microscopy of spherical droplets of 50 µM N^SARS-CoV-2^ and 1 µM polyU in 20 mM NaPi, pH 7.5. Fluorescently labeled RNA (green) partitioned into the droplets. Scale bar, 20 µm. **(D)** Increase in N^SARS-CoV-2^- and RNA-concentration inside of N^SARS-CoV-2^/polyU droplets in 20 mM NaPi, pH 7.5. Scale bars 3 µm. **(E)** Time-dependent change in diffusion of N^SARS-CoV-2^ inside polyU-induced droplets observed by FRAP. FRAP of freshly prepared droplets is shown in green, and after incubation for one hour in blue. Error bars represent the standard deviation for averaged six curves for each time point. Representative micrographs of a fresh droplet (top) and after incubation for one hour (bottom) before bleaching, after bleaching and at the end of recovery are displayed to the right.

Next, we monitored LLPS for different N^SARS-CoV-2^ and polyU concentrations. According to turbidity measurements, polyU-induced LLPS started at 5-10 µM N^SARS-CoV-2^ (Figure 1B and S1C). The polyU concentration at which maximum turbidity was observed shifted to higher polyU concentrations with increasing protein concentration (Figure 1B and S1C). Calculation of the N^SARS-CoV-2^ and polyU charge showed that LLPS is strongest when charge neutralization occurs (Figure 1B). At a given protein concentration, the turbidity increased with increasing polyU concentration, reached a maximum and then rapidly decreased to its starting value (Figure 1B), i.e. charge-matching RNA concentrations enable phase separation, but high RNA/protein ratios prevent LLPS of N^SARS-CoV-2^. The RNA-induced LLPS behavior of N^SARS-CoV-2^ is in agreement with the known properties of RNA-induced phase separation of prion-like RNA binding proteins (Maharana et al., 2018).

### Time-Dependent Transformation of N^SARS-CoV-2^/RNA-Droplets

A characteristic property of LLPS is the liquid-like nature of phase-separated compartments. We photobleached N^SARS-CoV-2^ inside of N^SARS-CoV-2^/polyU droplets and observed rapid recovery of fluorescence (Figure 1E). Exponential fitting of the recovery resulted in a diffusion coefficient of 0.198 ± 0.002 µm^2^/s. We then waited one hour and repeated fluorescence recovery after photobleaching (FRAP). At this later time point, the fluorescence recovery was best described by a bi-exponential fit consisting of two components with diffusion coefficients of 0.268 ± 0.063 µm^2^/s and 0.054 ± 0.020 µm^2^/s (Figure 1E and S1D). In addition, the fluorescence did not fully recover (Figure 1E), i.e. ∼60 % of N^SARS-CoV-2^ had transformed into an immobile species. The analysis shows that N^SARS-CoV-2^/polyU droplets change their material properties in a time-dependent manner. Because a major activity of N^SARS-CoV-2^ is to encapsulate RNA, the immobile fraction observed by FRAP might represent early stages of nucleocapsid assembly. Successful nucleocapsid formation, however, also depends on the specific sequence and secondary structure of viral RNA (Zhou and Routh, 2020) and is therefore not expected for polyU.

### SARS-CoV-2 Nucleocapsid Protein Associates with Stress Granules

Stress granules (SGs) are cytoplasmic RNA/protein granules, which form through LLPS and are modulated by corona- and other viruses to maximize replication efficiency (White and Lloyd, 2012). SARS-CoV-2 protein interaction mapping indicated that N^SARS-CoV-2^ binds the SG protein G3BP1 (Gordon et al., 2020). To investigate if N^SARS-CoV-2^ associates with SGs, we used a previously established SG-colocalization assay (Hofweber et al., 2018). SGs were induced in HeLa cells by arsenite followed by permeabilization of the plasma membrane by digitonin. Subsequently, soluble cytosolic factors were washed out and fluorescently labeled N^SARS-CoV-2^ was added together with Alexa Fluor 594-coupled antibody against the SG marker G3BP1. Laser scanning confocal microscopy showed that arsenite induced formation of SGs that stained for G3BP1 (Figure 2A). N^SARS-CoV-2^ colocalized with G3BP1-positive SGs (Figure 2A; Movie S1). FRAP of SG-associated N^SARS-CoV-2^ suggested the presence of three N^SARS-CoV-2^ populations (Figure 2B): a very mobile with rapid fluorescence recovery, a slower diffusing component and an immobile fraction, which does not recover its fluorescence after photobleaching (Figure 2C). Because SGs consist of a rigid core and a dynamic shell (Jain et al., 2016), we attribute the different N^SARS-CoV-2^ diffusion properties to the localization of N^SARS-CoV-2^ to different sub-structures of SGs. In agreement with our findings for N^SARS-CoV-2^, the N protein of SARS-CoV, the causative agent of the SARS epidemic in 2002/2003, translocates to SGs in stressed SARS-CoV-infected cells (Peng et al., 2008).

**Figure 2.**
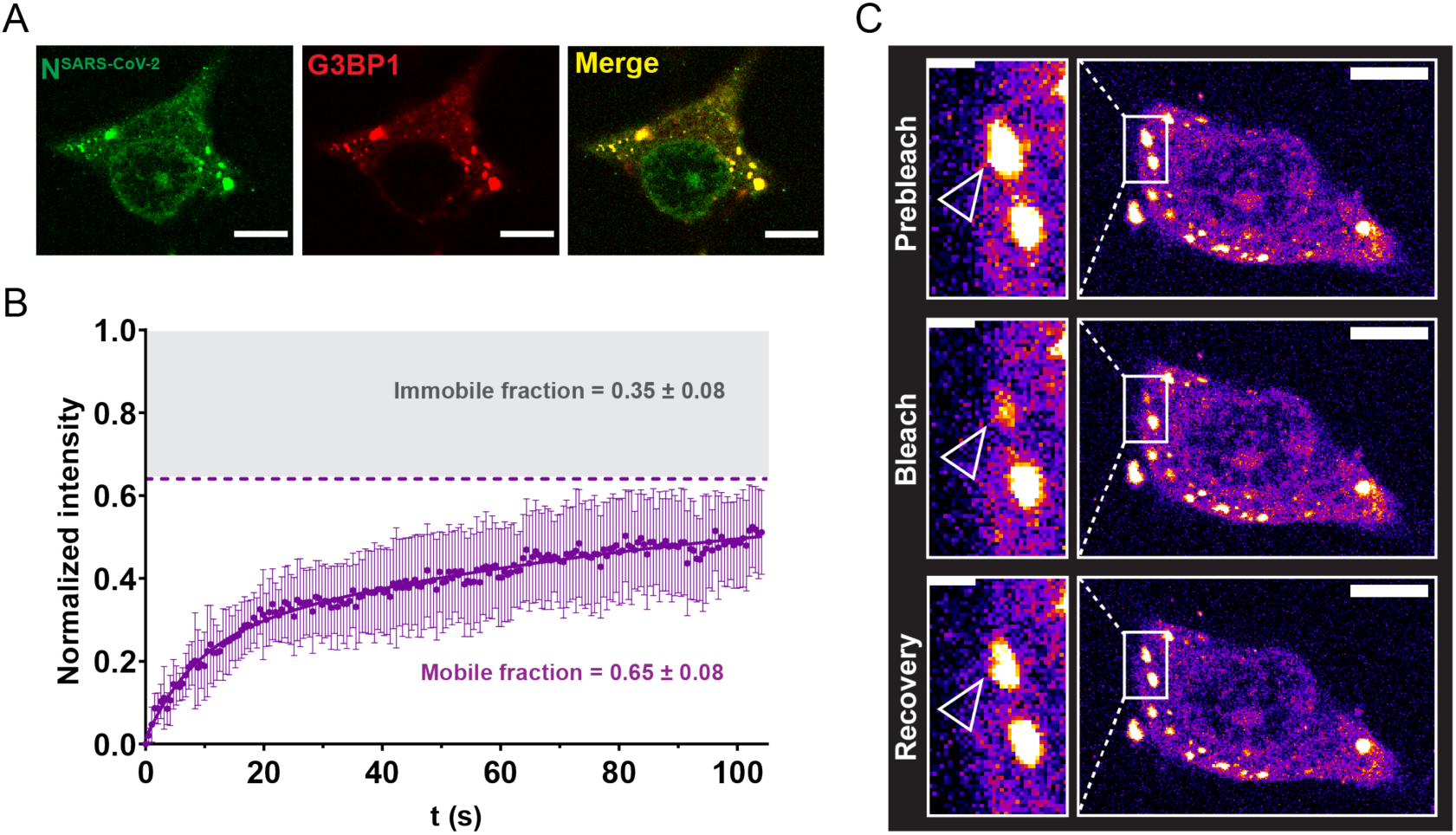
Nucleocapsid protein of SARS-CoV-2 associates with stress granules. **(A)** Alexa Fluor 488 labeled N^SARS-CoV-2^ (green) colocalizes with the stress granule marker G3BP1 (red) in arsenite-treated digitonin-permeabilized HeLa cells. **(B)** Fluorescence recovery after photobleaching (FRAP) curve fitted with bi-exponential fit suggests existence of a fast diffusing population of molecules (diffusion coefficient D_1_=0.166 ± 0.021 µm^2^/s), which together with a slowly recovering population (D_2_=0.014 ± 0.004 µm^2^/s) comprises ∼65% of the bleached spot (mobile fraction). The remaining 35% are immobile and do not recover its fluorescence after photobleaching. **(C)** Corresponding confocal microscopy pictures of the representative FRAP of N^SARS-CoV-2^ associated with stress granules in HeLa cells. Insert shows the bleached stress granule marked with an arrow. Scale bar 10 µm in (A) and (C), 2 µm in inset in (C). See also Movie S1.

### RNA-Interaction of the Mutation-Prone SR-Region

RNA viruses have enormously high mutation rates enhancing virulence and evolvability. On June 7^th^ 2020, already 42176 SARS-CoV-2 sequences were deposited (https://www.gisaid.org). Analysis of the corresponding N proteins showed that the mutations are most frequent in the SR-region of N^SARS-CoV-2^ (Figure 3A), which is conserved among human coronaviruses (Figure S2). SR-regions bind both RNA and proteins (Wegener and Muller-McNicoll, 2019). To gain insight into the molecular properties of the SR-region of N^SARS-CoV-2^ and its interaction with RNA, we combined NMR spectroscopy with molecular dynamics (MD) simulations. We particularly focused on the region from A182 to S197, because it contains 9 serine and 4 arginine residues, i.e. the highest density of SR-motifs in N^SARS-CoV-2^ (Figure S2). Chemical shift analysis showed that residues A182-S197 are very dynamic with a small propensity of α-helical structure next to R189 (Figure 3B-C).

**Figure 3.**
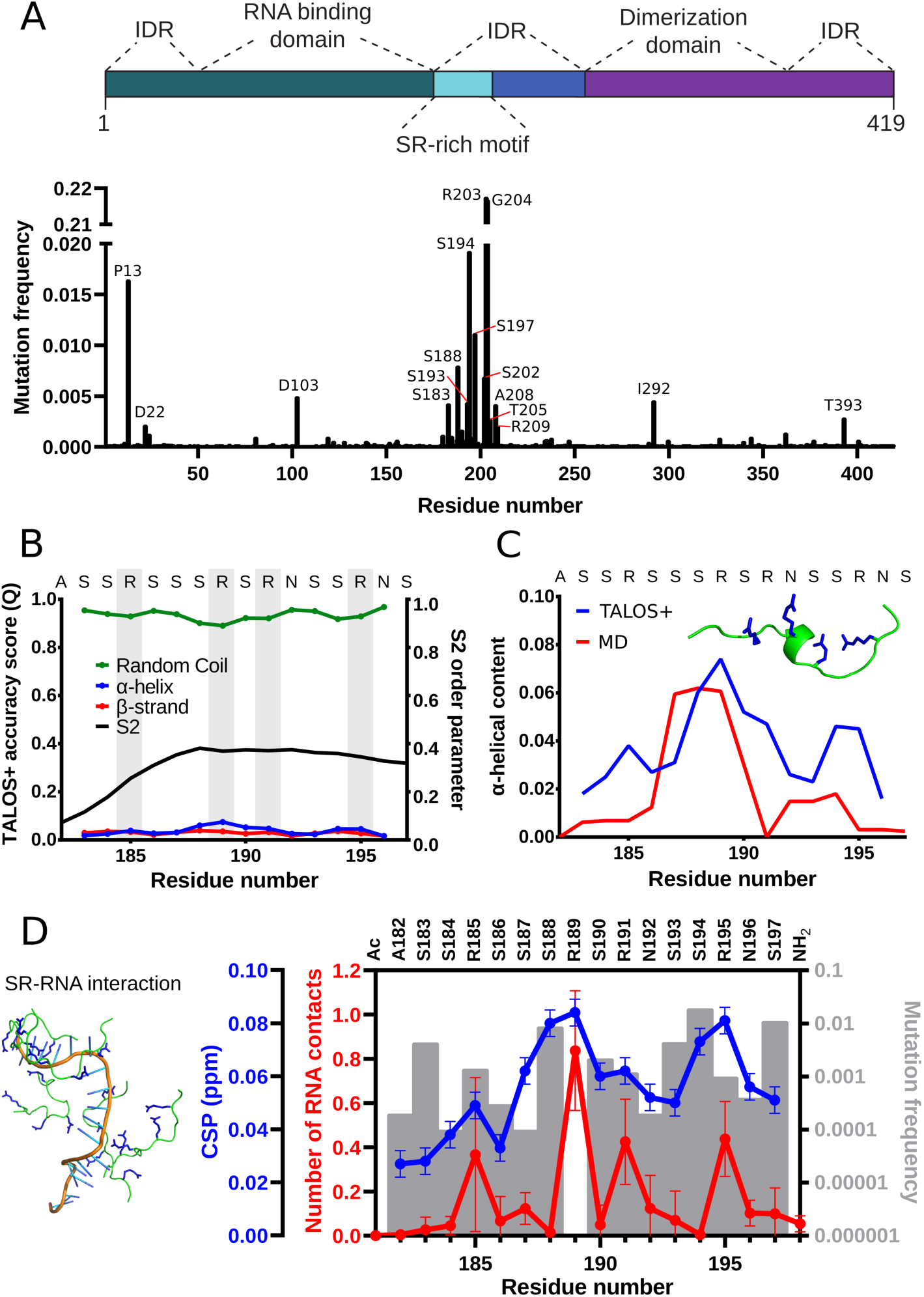
Atomic details of the RNA-interaction of the mutation-prone SR-region. **(A)** Frequency of mutations in the nucleocapsid protein in 42176 SARS-CoV-2 sequences from the China National Center for Bioinformation. Residues with more than 0.0025 frequency are labeled. Domain organization of N^SARS-CoV-2^ on top. **(B)** NMR-based analysis of the structure of the high-density SR-stretch (residues A182-S197) of N^SARS-CoV-2^. Secondary structure derived from chemical shifts using TALOS+ (Shen et al., 2009) is represented together with the S^2^ order parameter. Arginines are highlighted in grey. See also Figure S2. **(C)** Comparison between the α-helical propensity derived from NMR data (blue) and MD simulations (red). One conformer with α-helical content from the simulation is shown inside the graph. **(D)** Comparison between the number of peptide-polyU contacts per peptide in the MD simulations (red; average and standard deviation of five simulations), the NMR chemical shift perturbation (CSP) of the peptide-polyU titration at 1500 nM of polyU (blue), and the frequency of mutations per residue (grey bars). At the left, a MD snapshot of five SR-peptides with the 20 base length polyU is shown. See also Figures S3 and S4.

Next we investigated the conformational properties of A182-S197 with MD simulations. For the simulations, we used the state-of-the art force field/water model that accurately reproduced NMR parameters of the intrinsically disordered protein α-synuclein (Pietrek et al., 2020). In agreement with the NMR analysis, residues next to R189 populate transient α-helical structure (Figure 3C). We then performed calculations in the presence of polyU. A large number of intermolecular contacts between the arginine residues and the RNA phosphate groups were observed. Most intermolecular contacts were present for R189 (Figure 3D, red). In addition, R189 was most sensitive to the addition of polyU as observed by NMR spectroscopy (Figure 3D, blue; Figure S3A). R189 is the only residue in the region from A182-S197 that is not mutated in 42176 SARS-CoV-2 sequences (Figure 3D, grey bars), in agreement with its functional relevance.

### SR-Phosphorylation Modulates RNA-Induced Phase Separation

Phosphorylation of SR-domains provides functional specificity and adjustability to ribonucleoprotein formation (Wegener and Muller-McNicoll, 2019) and impairs binding of SR-domains of pre-mRNA splicing factors to protein hydrogel droplets (Kwon et al., 2014). To gain insight into the impact of phosphorylation of the SR-region of N^SARS-CoV-2^ on its RNA binding, we performed MD simulations of the high-density SR-stretch carrying phosphate groups at different serine residues (Figure S4A). Even when only a single serine, S188, was phosphorylated, the number of intra- and intermolecular peptide/peptide contacts increased (Figure S4B-C). Multi-site phosphorylation further raised the number of contacts (Figure S4B-C), and the intermolecular peptide/peptide contacts reached a maximum when three serines were phosphorylated (Figure S4C), i.e. when the overall charge is around zero. The phosphorylation-induced increase in intra- and intermolecular contacts is predominantly due to the formation of salt bridges between the phosphate groups and arginine side chains (Figure S4B-C). Because of the dense network of intra- and interpeptide salt bridges, contact formation with RNA – either polyU or a structured RNA derived from the viral genome of SARS-CoV-2 – was strongly attenuated upon phosphorylation (Figure 4A and S5).

**Figure 4.**
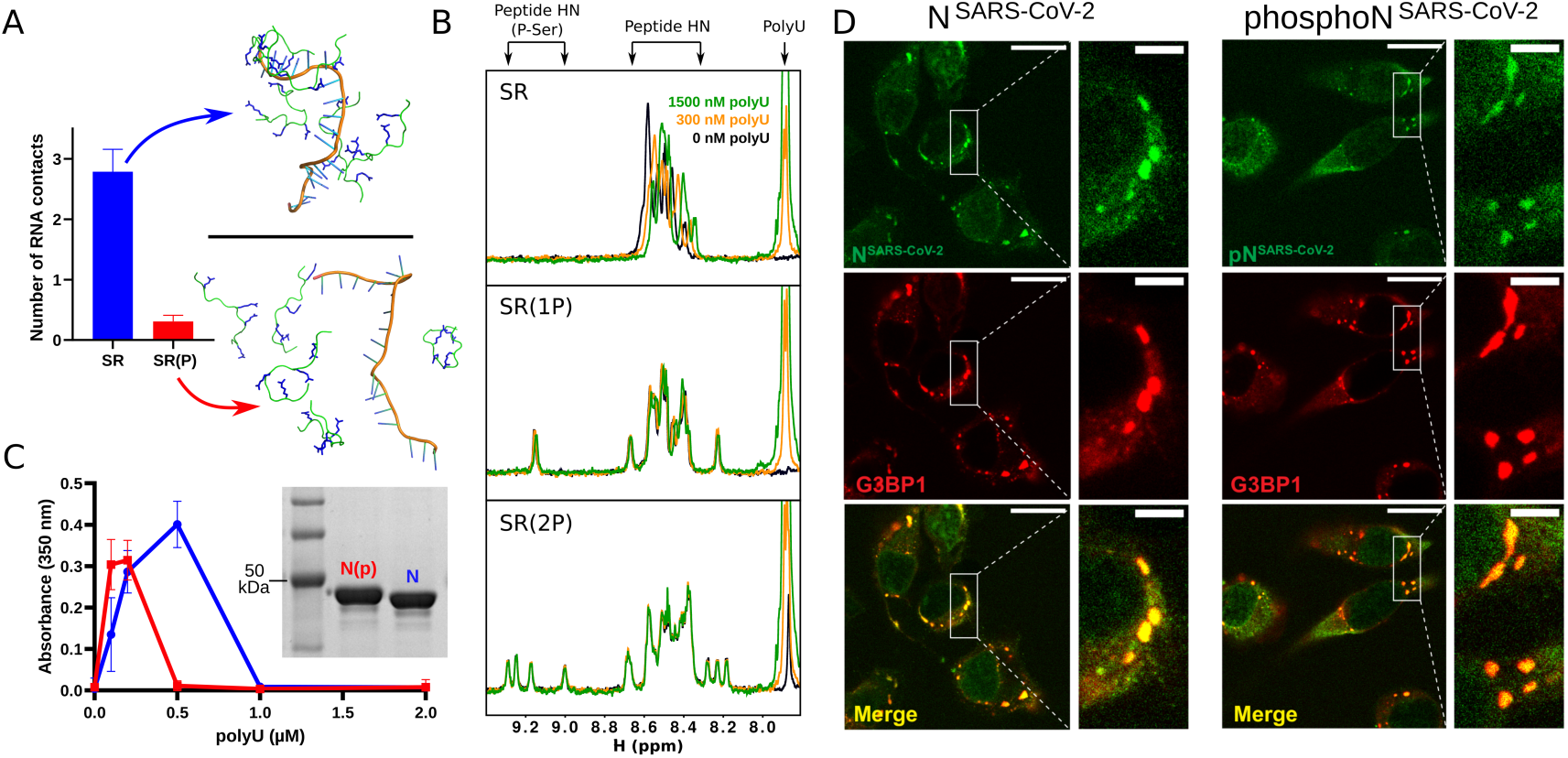
SR-phosphorylation modulates RNA-induced condensation of the nucleocapsid protein. **(A)** MD simulations of SR/polyU-interaction. The number of contacts per peptide with a 20 base length polyU is shown in the top graph for the non-phosphorylated and fully phosphorylated SR-peptide as the average and standard deviation of five independent simulations. Snapshots of both peptides are shown. See also Figure S5. **(B)** 1H 1D experiments of the SR-peptide (1766.8 Da), SPRK1 single-phosphorylated peptide (SR(1P); 1846.9 Da) and SPRK1 double-phosphorylated peptide (SR(2P); 1926.9 Da) at three different concentrations of polyU (0, 300 and 1500 nM). The spectral regions, in which the signals of polyU, the backbone H^N^s of unmodified residues and phosphorylated serines are located, are marked. The positive charges of the SR-peptide are compensated by the polyU negative charges at around 300 nM. **(C)** Turbidity at 350 nm of solutions of non-phosphorylated (blue) and SRPK1-phosphorylated (red) N^SARS-CoV-2^ in 20 mM NaPi, pH 7.5, at 30 μM protein concentration and increasing concentrations of polyU. Average values from three independent measurements are shown. Error bars, std. An SDS-Page gel (insert) displays a band shift due to SRPK1-phosphorylation. **(D)** Association of unmodified (N^SARS-CoV-2^, left panels) and phosphorylated (phosphoN^SARS-CoV-2^, right panels) nucleocapsid protein of SARS-CoV-2 with stress granules in HeLa cells. Scale bar 20 µm, 5 µm in inset.

To validate the results from MD simulation, we phosphorylated the SR-peptide *in vitro* using the serine/arginine protein kinase 1 (SRPK1) and performed NMR spectroscopy. SRPK1 phosphorylates SR-motifs and is involved in a wide spectrum of cellular activities including the regulation of viral genome replication (Fukuhara et al., 2006; Wegener and Muller-McNicoll, 2019). SRPK1-phosphorylation resulted in two species, a single phosphorylation at S188 (Figure 4B, middle) and a di-phosphorylated state, which is heterogeneously phosphorylated at four different serines (Figure 4B, bottom). NMR titrations showed that the unmodified SR-peptide strongly interacts with polyU, but not when it is phosphorylated at S188 (Figure 4B and S3B). LLPS experiments further demonstrated that phosphorylation of full-length N^SARS-CoV-2^ by SRPK1 changes its RNA-induced phase separation behavior (Figure 4C). The maximum of RNA-induced turbidity was shifted to lower polyU-concentrations for SRPK1-phosphorylated N^SARS-CoV-2^ (Figure 4C). In addition, co-localization of SRPK1-phosphorylated N^SARS-CoV-2^ with stress granules was slightly attenuated (Figure 4D). The combined data show that the SR-region is important for RNA-induced phase separation of N^SARS-CoV-2^, but multiple domains of N^SARS-CoV-2^ contribute to its translocation to stress granules. Phosphorylation and mutation in different regions of N^SARS-CoV-2^ (Figure 3A) are therefore poised to modulate viral replication.

### Nucleocapsid Protein LLPS Concentrates Components of the SARS-CoV-2 Replication Machinery

LLPS provides a cooperative mechanism to locally increase protein and RNA concentrations^11,12^. In addition, protein/RNA condensates can recruit additional proteins to promote reactions. To investigate if the RNA-dependent RNA polymerase (RdRp; Figure 5A) concentrates within N^SARS-CoV-2^/RNA droplets, we recombinantly prepared the non-structural protein (nsp) 12 of SARS-CoV-2, together with the accessory sub-units nsp7 and nsp8, which are required for transcription (Hillen et al., 2020). First we used nsp12, in order to investigate if the catalytic component of RdRp is recruited to N^SARS-CoV-2^/polyU droplets. Fluorescence microscopy revealed strong nsp12 fluorescence inside the droplets (Figure 5B).

**Figure 5.**
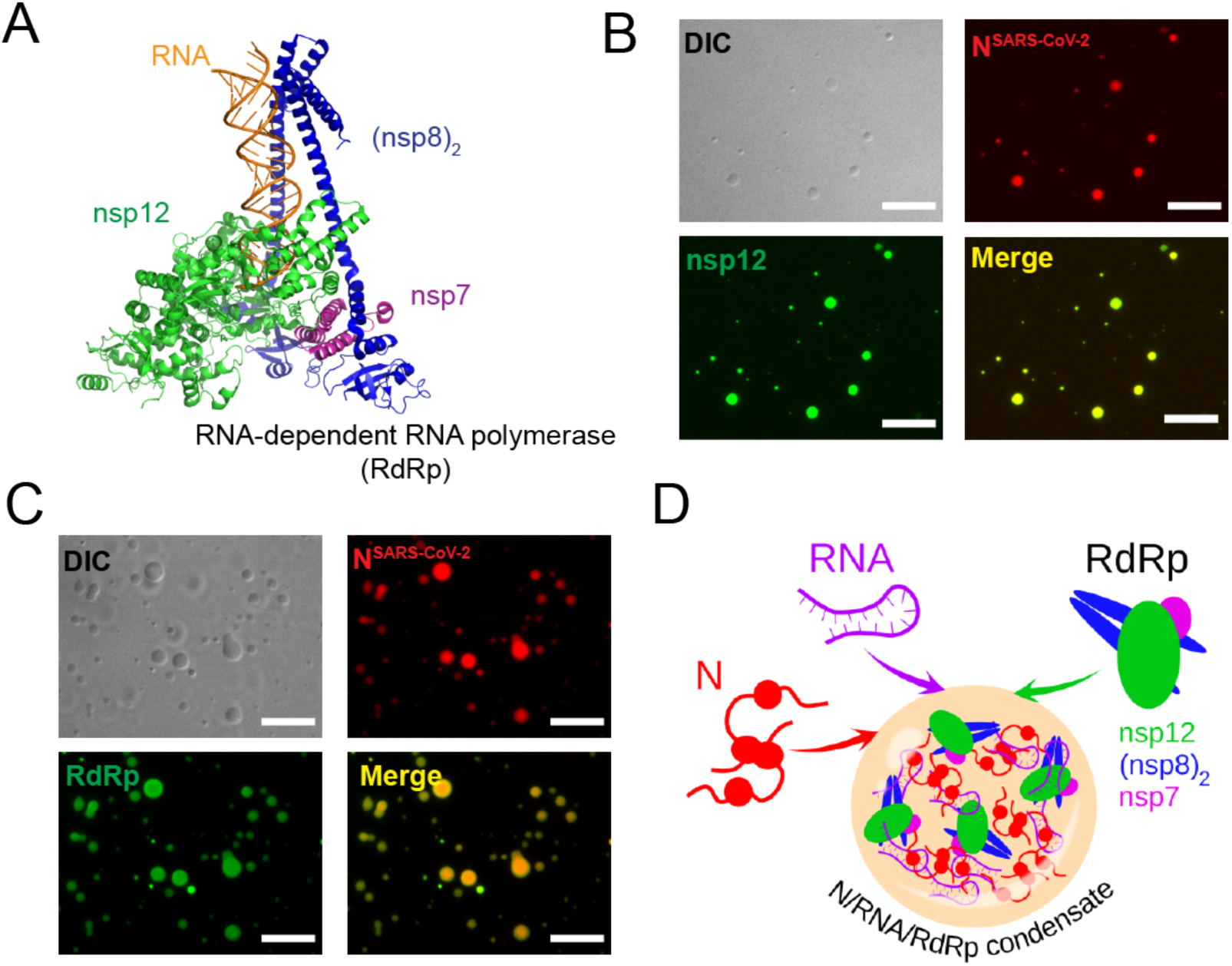
The RNA-dependent RNA polymerase complex of SARS-CoV-2 concentrates in RNA/nucleocapsid protein-droplets. **(A)** Structure of the RNA-bound RdRp-complex formed by the SARS-CoV-2 non-structural proteins nsp12 (green), nsp7 (magenta) and nsp8 (blue) in 1:1:2 stoichiometry (PDB code: 6YYT) (Hillen et al., 2020). **(B)** Fluorescence and DIC microscopy show recruitment of Alexa Fluor 488 labeled nsp12 (green), the catalytic subunit of the RdRp-complex, into N^SARS-CoV-2^/polyU droplets (N^SARS-CoV-^ 2 in red). **(C)** Active SARS-CoV-2 RdRp-complex bound to a fluorescein-labeled minimal RNA hairpin concentrates inside of N^SARS-CoV-2^/polyU droplets. In (B) and (C) droplets of 50 µM N^SARS-CoV-2^ and 1 µM polyU were prepared in 20 mM NaPi, pH 7.5, and visualized by addition of a small amount of Alexa Fluor 594 labeled N^SARS-CoV-2^ protein. Scale bars are 10 µm in (B) and (C). See also Figure S6. **(D)** Schematic representation of the LLPS-based formation of N/RNA/RdRp-condensates as protein/RNA-dense sites for viral transcription.

Next, nsp12, nsp7 and nsp8 were reconstituted in a 1:1:2 stoichiometry together with a RNA template-product duplex, which carried fluorescein at the 5’ end (Hillen et al., 2020). The RdRp/RNA-complex was added to preformed N^SARS-CoV-2^/polyU droplets into which the RdRp/RNA-complex was recruited (Figure 5C). High local concentrations of RdRp and N^SARS-CoV-2^ were reached (Figure S6). The data suggest that RNA-driven condensation of N^SARS-CoV-2^ provides a mechanism for bringing together components of the viral replication machinery (Figure 5D). In agreement with this proposed mechanism, N protein of SARS-CoV colocalizes intracellularly with replicase components (Stertz et al., 2007).

## DISCUSSION

Our study shows that the nucleocapsid protein of the SARS-CoV-2 virus undergoes RNA-induced liquid-liquid phase separation. Although nucleocapsid assembly can occur outside of liquid-like compartments, the rate of assembly is increased when the nucleocapsid protein is concentrated through LLPS (Guseva et al., 2020). In addition, N^SARS-CoV-2^ interacts with human ribonucleoproteins (Gordon et al., 2020), which are found in several LLPS-driven cytosolic protein/RNA granules, indicating that N^SARS-CoV-2^ modulates protein/RNA granule formation in order to maximize viral replication. In agreement with such activity, we showed that N^SARS-CoV-2^ translocates to stress granules in stressed cells.

We demonstrate that N^SARS-CoV-2^ LLPS promotes cooperative association of the RNA-dependent RNA polymerase complex with RNA. This suggests that SARS-CoV-2 uses LLPS-based mechanisms similar to transcription hubs in cellular nuclei (Boehning et al., 2018; Cramer, 2019) to enable high initiation and elongation rates during viral transcription. Because the replication machinery of coronaviruses is membrane-associated (Sola et al., 2015), it will furthermore be interesting to investigate if the SARS-CoV-2 glycoprotein M, which binds nucleocapsid protein (McBride et al., 2014), causes tethering of N^SARS-CoV-2^/RdRp/RNA-condensates to host cell membranes.

Taken together the data suggest that inhibition of the RNA-induced phase separation of the nucleocapsid protein of SARS-CoV-2 provides a viable and novel strategy for the design of therapeutics to treat Covid-19.

## METHOD DETAILS

### Materials

Alexa Fluor 594 conjugated anti G3BP1 antibody was purchased from Santa Cruz Biotechnology (sc-365338 AF594). Digitonin was from Merck (CAS 11024-24-1). PolyU potassium salt (800 kDa) was purchased from Sigma-Aldrich (Taufkirchen, Germany). Recombinant nucleocapsid protein of SARS-CoV-2 (Catalog No: <u>Z03488- 1</u>) and SRPK1 kinase (Catalog No: PV4215) were purchased from GenScript and<u>ThermoFisher</u>, respectively. The SR peptide, comprising residues 182-197 of the nucleocapsid protein of SARS-CoV-2, was produced by Fmoc-solid-phase synthesis using an automated microwave peptide synthesizer (Liberty 1, CEM) with acetyl-and amide protection groups at the N- and C-termini, respectively, and further purified by reverse-phase high-performance liquid chromatography (LC). Peptide masses were determined by LC-MS using an Acquity Arc System (WATERS) with SQD2-mass-detector.

### Purification of nsp12, nsp7, nsp8 and formation of RdRp complex

SARS-CoV-2 nsp12 was expressed in insect cells and purified as described previously(Hillen et al., 2020). SARS-CoV-2 nsp7 and nsp8 were expressed in *E.coli* and purified as described previously(Hillen et al., 2020). An RNA scaffold for RdRp/RNA complex formation was annealed by mixing equimolar amounts of two RNA strands (5’-rUrUrUrUrCrArUrGrCrArUrCrGrCrGrUrArG rGrCrUrCrArUrArCrCrGrUrArUrUrGrArGrA -3’; 56-FAM/rUrCrUrCrArArUrArCrGrGrUrArUrGrArGrC CrUrArCrGrCrG-3’) (IDT Technologies) in annealing buffer (10 mM Na-HEPES pH 7.4, 50 mM NaCl) and heating to 75 °C, followed by step-wise cooling to 4 °C. For complex formation, 2.45 nmol of purified nsp12 was mixed with a 1.1-fold molar excess of RNA scaffold and 3-fold molar excess of each nsp8 and nsp7. After incubation at room temperature for 10 min, the complex was subjected to size exclusion chromatography on a Superdex 200 Increase 3.2/300 equilibrated with complex buffer (20 mM Na-HEPES pH 7.4, 100 mM NaCl, 1 mM MgCl_2_, 1 mM TCEP). Peak fractions with a volume of approx. 125 µL (absorbance at 280 nm of 2.7 AU, 10 mm path length) corresponding to a nucleic acid-rich high-molecular weight population (as judged by absorbance at 260 nm) were pooled and used for subsequent experiments.

### Turbidity measurements

Phase diagrams of non-phosphorylated and phosphorylated N^SARS-CoV-2^ at different concentrations were determined using a NanoDrop spectrophotometer (ThermoFisher Scientific, Invitrogen). Increasing concentrations of polyU (0 - 2 µM) were added immediately before the experiments, followed by thoroughly pipetting and measurement of turbidity at 350 nm UV-Vis. Averages turbidity values were derived from measurements of three independent, freshly prepared samples.

### Microscopy

For fluorescence microscopy, proteins were labeled using Alexa-fluor 488™(green) or Alexa-fluor 594™(red) microscale protein labeling kits (ThermoFisher Scientific, Invitrogen). Small amounts (∼0.3 µl) of fluorescently-labeled N^SARS-COV-2^ were premixed with unlabeled N^SARS-COV-2^ and diluted to 50 µM final concentration with NaPi 20 mM buffer, pH 7.5. PolyU was added to the mixture to reach a final concentration of 1 µM. 5 µl of the sample were subsequently loaded onto a slide and covered with a 18 mm coverslip. DIC and fluorescent micrographs were acquired on a Leica DM6B microscope with a 63× objective (water immersion) and processed using Fiji software (NIH). For RNA recruitment assays, fluorescently labeled RNA was premixed with 1 µM polyU and subsequently added to the mixture of fluorescently labeled/unlabeled N^SARS-COV-2^. For nsp12/RdRp/RNA recruitment assays, the fluorescently labeled component (FAM-labeled RNA scaffold, nsp12 or RdRp/RNA-complex) was premixed with 1 µM polyU, followed by addition to a mixture of fluorescently labeled/unlabeled N^SARS-COV-2^.

### Fluorescence recovery after photobleaching (FRAP)

Dynamics of N^SARS-CoV-2^ molecules in the phase-separated state were investigated by FRAP analysis. As described above, N^SARS-CoV-2^ phase separation was induced using 50 µM N^SARS-CoV-2^ and 1 µM polyU in 20 mM sodium phosphate buffer (NaPi), pH 7.5. Droplet movement was reduced through addition of 5 % dextran T500 (Pharmacosmos). The sample was spiked with a small amount of Alexa Fluor 488 lysine labeled N^SARS-CoV-2^, which was photobleached and followed during recovery. FRAP recordings were done on freshly formed droplets settled down on the glass slide (early time points; time of recording <15 min) and after one hour of sample incubation on the glass (late time points).

FRAP experiments were recorded on a Leica TCS SP8 confocal microscope using a 63x objective (oil immersion) and a 488 argon laser line. A circular region of ∼4 µm in diameter was chosen in a region of homogenous fluorescence away from the droplet boundary and bleached with five iterations of full laser power. Recovery was imaged at low laser intensity (5%). 100 frames were recorded with one frame per 523 ms. Pictures were analyzed in FIJI software (NIH) and FRAP recovery curves were calculated using standard methods. Briefly, fluorescence intensities were measured for prebleaching, bleached and reference ROI in order to calculate the diffusion coefficient from FRAP curves. The prebleaching ROI was a selected region in the droplet before bleaching; the bleached ROI corresponded to the bleached area while the reference ROI corresponded to an area which did not experience bleaching. The fluorescence intensity measured for each of the described ROIs was corrected by background substraction: a region where no fluorescence was detected was used to calculate the background. Thus, the FRAP recovery was calculated as:

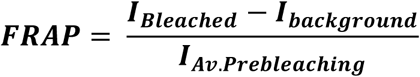

The value obtained was then corrected by multiplication with the acquisition bleaching correction factor (ABCF), which was calculated according to:

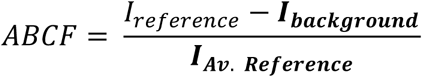

Finally, the curves were normalized according to:

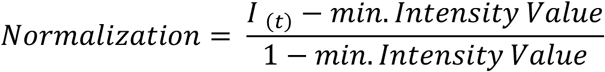

Values were averaged from six recordings for both early and late time points and the resulting FRAP curves ± standard deviation (std) were fitted for the early time points to a mono-exponential function:

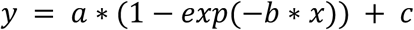

For the late time points, a bi-exponential function provided the best fitting:

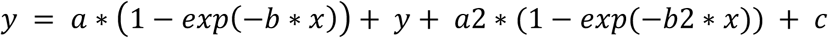

In both equations, b/b2 were used to calculate the half time 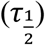 of recovery. The obtained value was then used to calculate the diffusion coefficient according to the equation:

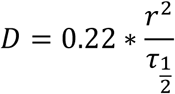

where *r* is the radius of the bleached area.

The mobile and immobile fractions were calculated using the parameters a and c derived from each fitting, according to the following equations:

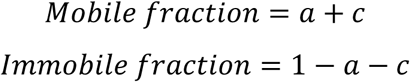

### Stress granule co-localization and FRAP

HeLa cells (DSMZ-German Collection of Microorganisms and Cell Cultures GmbH, ACC 57) were grown in an incubator at 37 °C in humidified atmosphere with 5 % CO_2_. One day before the stress granule co-localization assay and FRAP measurements cells were seeded in 96-well CELLview™slides (Greiner Bio-One) with glass bottom suitable for imaging so that on the day of the measurement cells reached ∼50% of confluence. Cell membranes were permeabilized by incubation in cell permeabilization buffer (20 mM HEPES-KOH pH 7.5, 120 mM KOAc, 5 mM Mg(OAc)_2_, 250 mM sucrose) supplemented with 60 µg/ml digitonin for 40 s. After washing cells four times with 100 µL of cell permeabilization buffer live recording was started on a Leica TCS SP8 confocal microscope using a 63x objective (oil immersion) and 488 argon laser or 561 laser lines. A mixture of 0.5 µm Alexa Fluor 488 lysine labeled N^SARS-CoV-2^ and 1:100 Alexa Fluor 594 conjugated G3BP1 antibody (Santa Cruz Biotechnology, sc-365338 AF594) in cell permeabilization buffer was added to the cells and movies were recorded with 512 x 512 pixel resolution at 1000 Hz speed and 1 s per frame for about 2-3 minutes. For FRAP measurements, individual stress granules were marked and photobleached with five iterations of full 488 argon laser power. Recovery was imaged at low laser intensity (8 %). 200 frames were recorded with one frame per 523 ms. Pictures were analyzed in FIJI software (NIH) and FRAP recovery curves ± standard deviation were calculated and fitted using bi-exponential fit as described above. Non-bleached stress granules were used as reference for calculations.

### In vitro phosphorylation

Phosphorylation of a stock of 850 µM SR-peptide (1766.8 Da) was performed by incubation with 0.15 µM SRPK1 kinase at 23 °C overnight in a buffer containing 4 mM ATP, 5 mM MgCl_2_, 1 mM DTT and 5 mM EGTA. Because of the intrinsically disordered nature of the peptide, inactivation of the kinase was achieved by incubation of the sample at 65 °C for 20 minutes, followed by centrifugation at 15,000 g for 30 minutes. Residual ATP, MgCl_2_ and EGTA were removed by HPLC followed by mass spectrometry. Phosphorylation of 100 µM unlabeled N^SARS-COV-2^ was performed by incubation with 0.5 µM of SRPK1 kinase. The reaction mixture was incubated at 23 °C overnight in a buffer containing 8 mM ATP, 5 mM MgCl_2_, 1 mM DTT and 5 mM of EGTA. Residual ATP, MgCl_2_ and EGTA were removed by 4 times buffer exchange using a Vivaspin 500.5 molecular weight cut-off (Sartorius, Göttingen). Samples were loaded onto a SDS-PAGE gel to confirm phosphorylation.

### Mutation frequency analysis

To examine the mutation frequency in SARS-CoV-2 sequences we used the database and resources from the China National Center for Bioinformation, 2019 Novel Coronavirus Resource (https://bigd.big.ac.cn/ncov?lang=en; downloaded June 7, 2020, with 42,176 genome sequences). The mutations between the genome positions 28274 and 29530 were analyzed to get the mutation frequency for each codon.

### NMR spectroscopy

One-dimensional (1D) ^1^H NMR experiments and two-dimensional (2D) ^1^H-^1^H TOCSY, NOESY and ^1^H-^15^N/^1^H-^13^C heteronuclear single quantum coherence (HSQC) experiments of the SR-peptide (residues A182-S197) of N^SARS-CoV-2^ were acquired at 5 °C on a Bruker 700 MHz spectrometer equipped with a triple-resonance 5 mm cryogenic probe. The peptide concentration was 4 mM for resonance assignment and 200 µM for the interaction analysis with polyU (800 kDa). Samples were in 50 mM NaP, 0.01 % NaN_3_ and 5 % D_2_O. Spectra were processed with TopSpin 3.6 (Bruker) and analyzed using Sparky (Lee et al., 2015). Secondary structure was analyzed subjecting experimental HA, HN, N, CA and CB chemical shifts to TALOS+ (Shen et al., 2009). The chemical shift perturbation (CSP) for the peptide residues is the one of the NH protons from the TOCSY experiment. The CSP error is based on the resolution of the spectra.

### Molecular dynamics simulations

Starting structures of the SR-peptides were built in the PyMOL Molecular Graphics System (Version 1.8.4.0, Schrödinger, LLC), those of the RNA molecules using the RNA modelling software SimRNA (Magnus et al., 2016). Initially, the different mixtures were equilibrated with 50,000 steps of energy minimization. To further equilibrate the system, 100 ps each of volume (NVT) and pressure (NPT) equilibration were performed without position restrains in order to have different starting points in each simulation. The MD simulations were carried out in GROMACS (version 2018.3) using the AMBER99SB-ILDN force field and the TIP3P water model at a temperature of 300 K, 1 bar of pressure and with a coupling time (ζT) of 0.1 ps. The mixtures were solvated in water with 150 mM NaCl, ensuring overall charge neutrality. The particle mesh Ewald algorithm was used for calculation of the electrostatic term, with a radius of 16 Å for the grid-spacing and Fast Fourier Transform. The cut-off algorithm was applied for the non-coulombic potential with a radius of 10 Å. The LINCS algorithm was used to contain bonds and angles. MD simulations were performed during 1 ns in 2 fs steps and saving the coordinates of the system every 10 ps. The force field parameters for the phosphorylated amino acids were taken from (Craft and Legge, 2005). The number of contacts and secondary structure over the simulation trajectory were analyzed using the PyMOL Molecular Graphics System (Version 1.8.4.0, Schrödinger, LLC). To get error bars, 5 repetitions were done for each simulation.

## SUPPLEMENTAL INFORMATION

Supplemental Information can be found online at

## ACKNOWLEDGMENTS

We thank Hauke S. Hillen, Goran Kokic, Lucas Farnung and Patrick Cramer for preparation of nsp12 and the RdRp complex, and Kerstin Overkamp for peptide synthesis and mass spectrometry. M.Z. was supported by the advanced grant ‘787679 - LLPS-NMR’ of the European Research Council and the German Science Foundation through SPP2191.

## AUTHOR CONTRIBUTIONS

A.S. conducted LLPS assays; A.I.d.O. performed bioinformatic analysis, NMR spectroscopy and MD simulations; M.R. performed FRAP and stress granule experiments; A.S., A.I.d.O., M.R. and M.Z. designed the project and wrote the paper.

## DECLARATION OF INTERESTS

The authors declare no competing interests.

## Notes

### Competing Interest Statement

The authors have declared no competing interest.

