## Supplementary material for "Nucleocapsid protein of SARS-CoV-2 phase separates into RNA-rich polymerase-containing condensates": savastano-SI.pdf

**Movie S1. Colocalization of $N^{SARS\text{-}CoV\text{-}2}$ and the stress granule marker G3BP1 in arsenite-induced stress granules in HeLa cells.** Scale bar 10 µm.

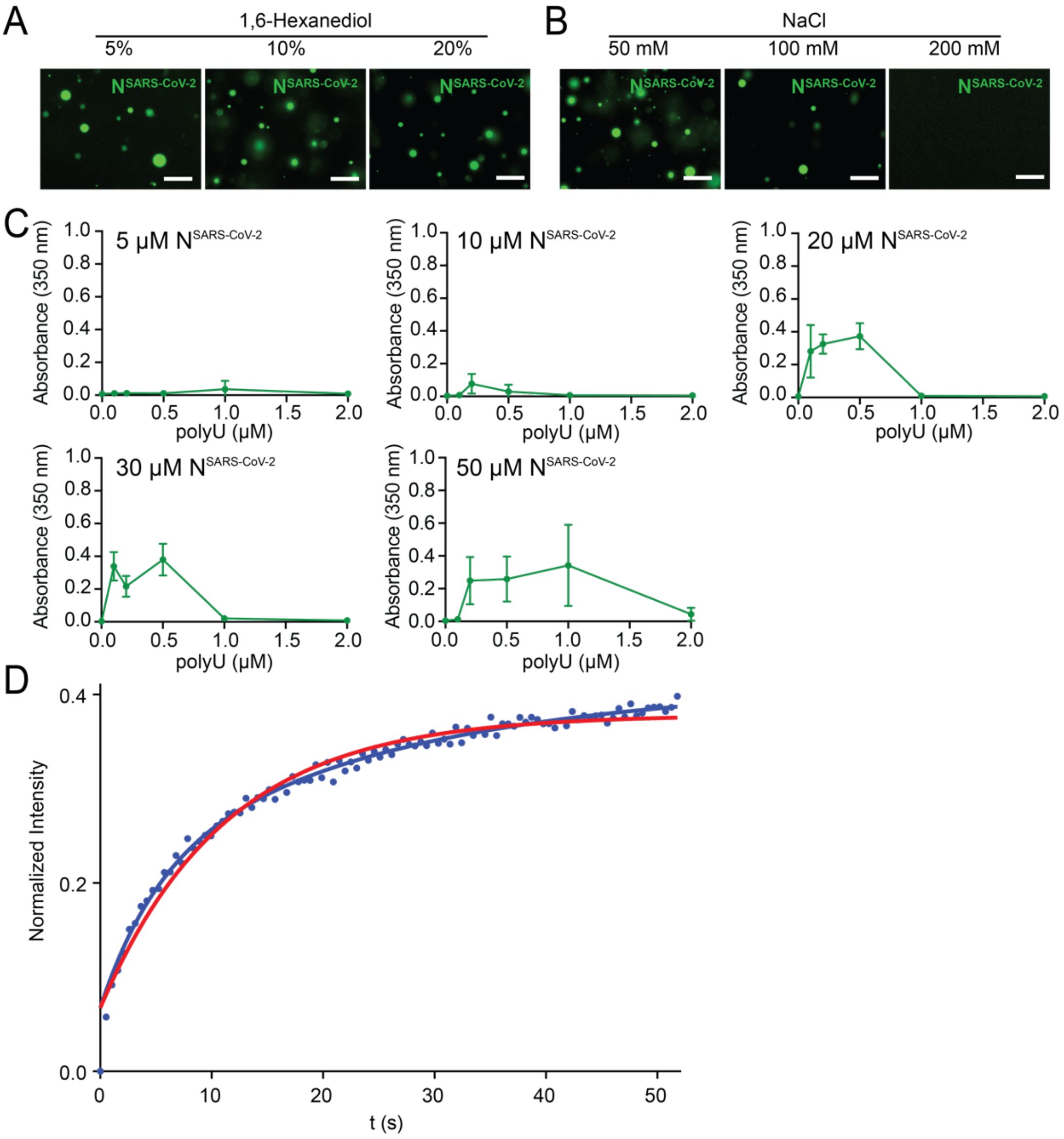


**Figure S1. RNA-induced LLPS of N$^{SARS\text{-}CoV\text{-}2}$**

(**A-B**) Fluorescence microscopy of droplets formed by 50 µM N$^{SARS\text{-}CoV\text{-}2}$ and 1 µM polyU in 20 mM NaPi, pH 7.5, at increasing concentrations of 1,6-hexanediol (A) or NaCl (B). Scale bars, 20 µm.

(**C**) Turbidity at 350 nm of solutions of N$^{SARS\text{-}CoV\text{-}2}$ in 20 mM NaPi, pH 7.5, at different protein concentrations (5-50 µM) and increasing concentrations of polyU. Average values from three independent measurements are shown and also displayed in Figure 1B. Error bars, std.

(**D**) Fit of a mono-exponential (red) and bi-exponential (blue) function to FRAP data obtained for N$^{SARS\text{-}CoV\text{-}2}$/polyU droplets incubated for one hour.

```
SARS-CoV-2     1 MSDNG-PQ-NQRNA-------PR-ITFGGPSDSTGSNQNGERSGARSKQ---RR-PQ-GL
SARS-CoV       1 MSDNG-PQSNQRSA-------PR-ITFGGPTDSTDNNQNGGRNGARPKQ---RR-PQ-GL
MERS-Cov       1 MASPA--------A-------PRAVSFADNNDITNTNLSRGR-GRNP------K-PR-AA
HCoV-HKU1      1 MSYTPGHYAGSRSSSGNRSGILKKTSWADQSERNYQTFNRGR-KTQPKFTVSTQ-PQ-GN
HCoV-OC43      1 MSFTPGKQSSSRASSGNRS-VNGILKWADQSDQFRNVQTRGR-RAQPKQTATSQQPSGGN
HCoV-NL63      1 MASV---------------------NWADDRAA--------R-KK---------------
HCoV-299E      1 MATV---------------------KWADASEPQ-----RGR-QG---------------

SARS-CoV-2    46 PNNTASWFTALT-QHGK-EDLKFPRGQGVPINTNSSPDDQIGYYRRATR-RIRGGDGKMK
SARS-CoV      47 PNNTASWFTALT-QHGK-EELRFPRGQGVPINTNSGPDDQIGYYRRATR-RVRGGDGKMK
MERS-Cov      37 PNNTVSWYTGLT-QHGK-VPLTFPPGQGVPLNANSTPAQNAGYWRRQDR-KINTGNG-IK
HCoV-HKU1     58 TIPHYSWFSGIT-QFQKGRDFKFSDGQGVPIAFGVPPSEAKGYWYRHSRRSFKTADGQQK
HCoV-OC43     59 VVPYYSWFSGIT-QFQKGKEFEFAEGQGVPIAPGVPATEAKGYWYRHNRRSFKTADGNQR
HCoV-NL63     16 -FPPPSFYMPLLVSSDK-APYRVIPRNLVPIGKGN-KDEQIGYWNVQER--WRMRRGQRV
HCoV-299E     19 -RIPYSLYSPLL-VDSE-QPWKVIPRNLVPINKKD-KNKLIGYWNVQKR--FRTRKGKRV

SARS-CoV-2   103 DLSPRWYFYYLGTGPEAGLPYGANKDGIIWVATEGAL-NTPKDHIGTRNPANNAAIVLQL
SARS-CoV     104 ELSPRWYFYYLGTGPEASLPYGANKEGIVWVATEGAL-NTPKDHIGTRNPNNNAATVLQL
MERS-Cov      93 QLAPRWYFYYTGTGPEAALPFRAVKDGIVWVHEDGAT-DAPST-FGTRNPNNDSAIVTQF
HCoV-HKU1    117 QLLPRWYFYYLGTGPYANASYGESLEGVFWVANHQADTSTPSD-VSSRDPTTQEAIPTRF
HCoV-OC43    118 QLLPRWYFYYLGTGPHAKDQYGTDIDGVYWVASNQADVNTPAD-IVDRDPSSDEAIPTRF
HCoV-NL63     71 DLPPKVHFYYLGTGPHKDLKFRQRSDGVVWVAKEGAK-TVNTS-LGNRKRNQKPLEP-KF
HCoV-299E     73 DLSPKLHFYYLGTGPHKDAKFRERVEGVVWVAVDGAK-TEPTG-YGVRRKNSEPEIP-HF

SARS-CoV-2   162 PQGTTLPKGFYA-EGSRGGSQASSRSSSRS--RNSSRNSTPG-SSRGTSPARMAGNGG--
SARS-CoV     163 PQGTTLPKGFYA-EGSRGGSQASSRSSSRS--RGNSRNSTPG-SSRGNSPARMASGGG--
MERS-Cov     151 APGTKLPKNFHI-EGTGGNSQSSSRASSLS--RNSSRSSSQG-SRSGNS-TRGTSPGP--
HCoV-HKU1    176 PPGTILPQGYYV-EGS-GRSASNSRPGSRS----QSRGPNNRSLSRSNSNFRHSDSIV--
HCoV-OC43    177 PPGTVLPQGYYI-EGS-GRSAPNSRSTSRT----SSRASSAGSRSRANSGNRTPTSGV--
HCoV-NL63    128 S--IALPPELSVVEFE-DRSNNSSRASSRSSTRNNSRDSSRS-TSRQQSRTRSDSNQSSS
HCoV-299E    130 N--QKLPNGVTVVE-E-PDSRAPSRSQSRSQSRG--RGESKP-QSRNPSSDRNHNS--QD

SARS-CoV-2   216 ----DA--ALALLLLDRLNQLESKMSGK-GQQ-QQG-Q------------TV------TK
SARS-CoV     217 ----ET--ALALLLLDRLNQLESKVSGK-GQQ-QQG-Q------------TV------TK
MERS-Cov     204 SGIGAV--GGDLLYLDLLNRLQALESGK-VKQ-SQP-K------------VI------TK
HCoV-HKU1    228 ----K---------PDMADEIANLVLAKLGKD-SKP-Q------------QV------TK
HCoV-OC43    229 ----T---------PDMADQIASLVLAKLGKDATKP-K------------QV------TK
HCoV-NL63    184 DLVAAVTLALKNLGFDN--QSKSPSSSGTSTP-KKPNK----------------PLSQPR
HCoV-299E    181 DIMKAVAAALKSLGFDKP-QEKDKKSAKTGTP--KPSRNQSPASSQTSAKSLARSQSSET

SARS-CoV-2   249 KSAAE----ASKKPRQKRTATKA--YNVTQAFGRRGPEQTQGNFGDQELIRQGTDYKHWP
SARS-CoV     250 KSAAE----ASKKPRQKRTATKQ--YNVTQAFGRRGPEQTQGNFGDQDLIRQGTDYKHWP
MERS-Cov     241 KDAAA----AKNKMRHKRTSTKS--FNMVQAFGLRGPGDLQGNFGDLQLNKLGTEDPRWP
HCoV-HKU1    255 QNAKEIRHKILTKPRQKRTPNKH--CNVQQCFGKRGPS---QNFGNAEMLKLGTNDPQFP
HCoV-OC43    257 HTAKEVRQKILNKPRQKRSPNKQ--CTVQQCFGKRGPN---QNFGGGEMLKLGTSDPQFP
HCoV-NL63    225 ADKPS----QLKKPRWKRVPTRE--ENVIQCFGPRDFN---HNMGDSDLVQNGVDAKGFP
HCoV-299E    238 KEQKH----EMQKPRWKRQPNDDVTSNVTQCFGPRDLD---HNFGSAGVVANGVKAKGYP

SARS-CoV-2   303 QIAQFAPSASAFFGMSRIGMEVT-----------P-SGTWLTYTGAIKLDDKDPNFKDQV
SARS-CoV     304 QIAQFAPSASAFFGMSRIGMEVT-----------P-SGTWLTYHGAIKLDDKDPQFKDNV
MERS-Cov     295 QIAELAPTASAFMGMSQFKLTHQN----ND-DH-GNPVYFLRYSGAIKLDPKNPNYNKWL
HCoV-HKU1    310 ILAELAPTPGAFFFGSKLDLVKRDSEA---DSPVK-DVFELHYSGSIRFDSTLPGFETIM
HCoV-OC43    312 ILAELAPTAGAFFFGSRLELAKVQNLSGNPDEPQK-DVYELRYNGAIRFDSTLSGFETIM
HCoV-NL63    276 QLAELIPNQAALFFDSEVSTDEV-----------G-DNVQITYTYKMLVAKDNKNLPKFI
HCoV-299E    291 QFAELVPSTAAMLFDSHIVSKES-----------G-NTVVLTFTTRVTVPKDHPHLGKFL

SARS-CoV-2   351 ILLNKHIDAYKTFPP------TEPK--KDKKK-----KAD-ETQALPQRQKKQQTVT-LL
SARS-CoV     352 ILLNKHIDAYKTFPP------TEPK--KDKKK-----KTD-EAQPLPQRQKKQPTVT-LL
MERS-Cov     349 ELLEQNIDAYKTFPK------KEKKQKAPKEE-----STD-QMSEPPKEQRVQGSIT-QR
HCoV-HKU1    366 KVLEENLNAYVNSNQNTDSDSLSSKPQRKRGVKQLPEQFDSLNLSAGTQHISN-----DF
HCoV-OC43    371 KVLSENLNAYQQQDG---MMNMSPKPQRQRGHKNGQGENDNISVAVPKSRVQQNKSI-EL
HCoV-NL63    324 EQISAFTK---------------PSSIKEMQS-----QSS-HV---AQNTVLNASIPESK
HCoV-299E    339 EELNAFTREMQQHPL------LNPSALEFNPS-----QTS-PA--------------TAE
```

**Figure S2. Sequence alignment of the seven human coronaviruses**

The following sequences were used: SARS-CoV-2, *YP_009724397.2*; SARS-CoV, *NC_004718*; MERS-CoV, *NC_019843*; HCoV-HKU1, *NC_006577*; HCoV-OC43, *KF530099.1*; HCoV-299E, *NC_002645*; HCoV-NL63, *NC_005831*. Identical residues are marked in red, homologues residues in orange and yellow.

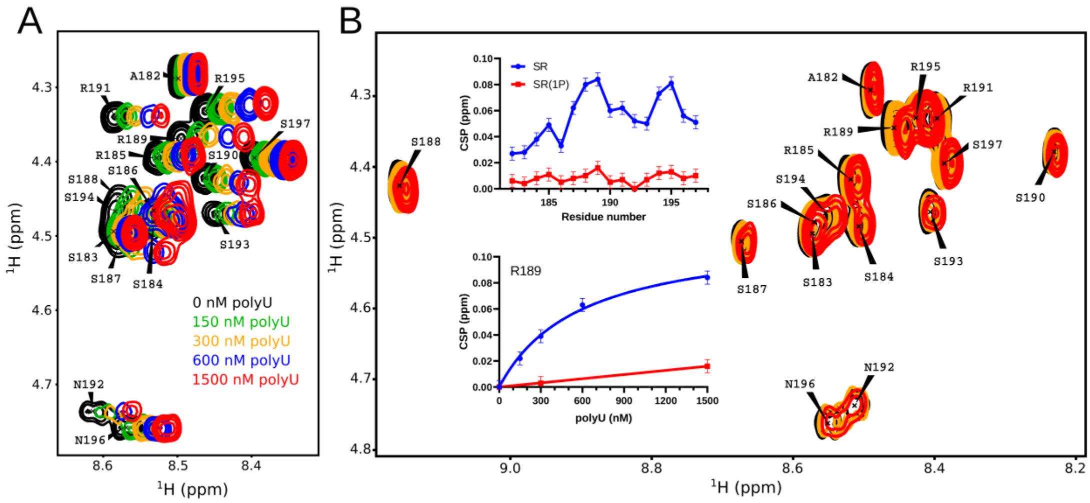


**Figure S3. NMR spectroscopy of the interaction of polyU with the SR-peptide comprising residues 182-197 of N$^{SARS\text{-}CoV\text{-}2}$**

(**A**) Selected regions from 2D TOCSY experiments of the SR-peptide at five different concentrations of polyU (0, 150, 300, 600 and 1500 nM). Resonance assignments are indicated.

(**B**) TOCSY spectra of the SPRK1 single-phosphorylated peptide at three different concentrations of polyU (0, 300 and 1500 nM). The positive charges of the SR-peptide are compensated by the negative charges of polyU at around 300 nM polyU. Spectral color code as in (A). The residue-specific chemical shift perturbation (CSP) observed for each peptide at 1500 nM of polyU are shown in the top plot. The change in R189 CSP with increasing polyU concentration is shown in the bottom plot. Data for the non-phosphorylated SR-peptide are shown in blue, for the SPRK1 single-phosphorylated peptide in red. The CSP error is based on the resolution of the spectra.

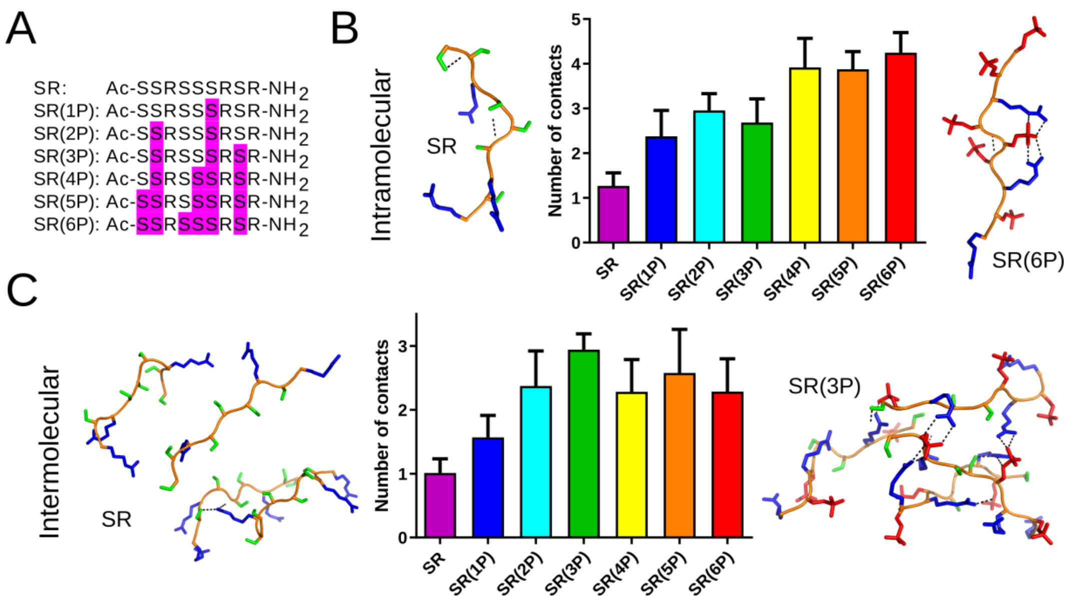


**Figure S4. Intra- and intermolecular interactions observed in MD simulations of SR-peptides comprising residues 183-191 of $N^{SARS\text{-}CoV\text{-}2}$**

(**A**) Amino acid sequences of the SR-peptides with phosphorylated serines highlighted in magenta.

(**B**) Number of intramolecular polar contacts per peptide for each SR-peptide displayed as the average and standard deviation of five independent simulations. Snapshots of the non-phosphorylated (left) and fully-phosphorylated SR-peptide (right) are shown.

(**C**) Number of intermolecular polar contacts per peptide for each SR-peptide displayed as the average and standard deviation of five independent simulations. MD simulations were performed with four identical peptides in the water box. Snapshots of non-phosphorylated (left) and 3-times phosphorylated SR-peptides (right). Salt bridges between the phosphate groups (red) and the arginine side-chains (blue) are marked by dashed lines.

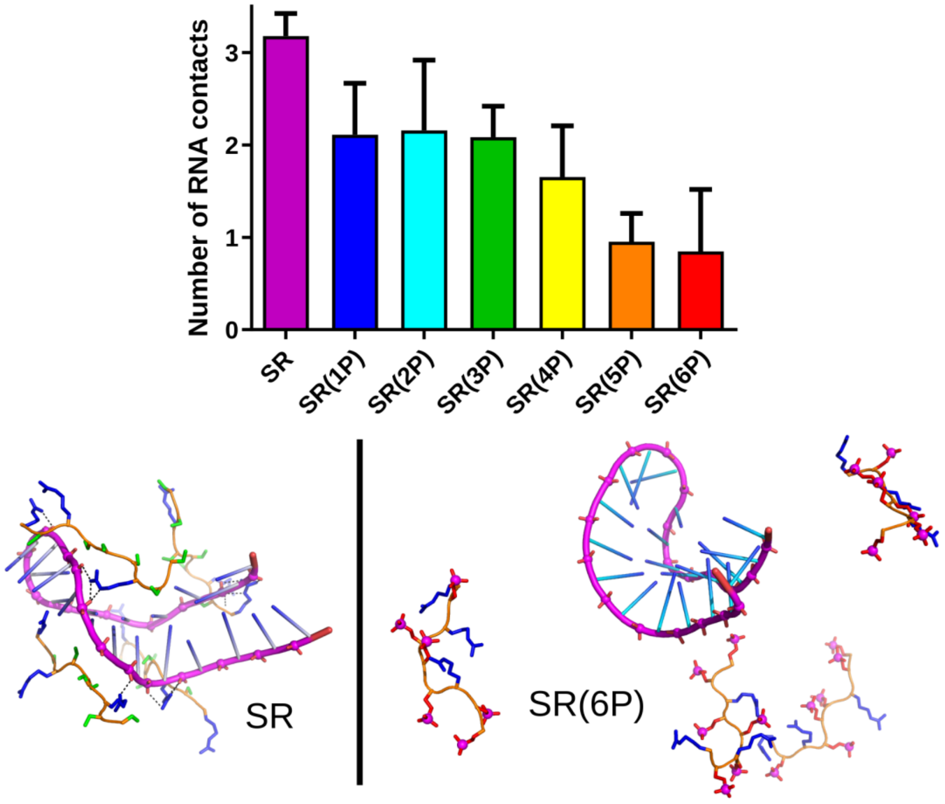


**Figure S5. RNA-interactions observed in MD simulations of SR-peptides comprising residues 183-191 of N$^{\text{SARS-CoV-2}}$**

The number of polar contacts of the different SR-peptides (see Fig. S4A) with a structured RNA derived from the viral genome of SARS-CoV-2 are displayed as the average and standard deviation of five independent simulations. MD snapshots of the RNA-interaction of the non-phosphorylated (left) and fully-phosphorylated (right) peptide are shown below. Arginine side chains in blue.

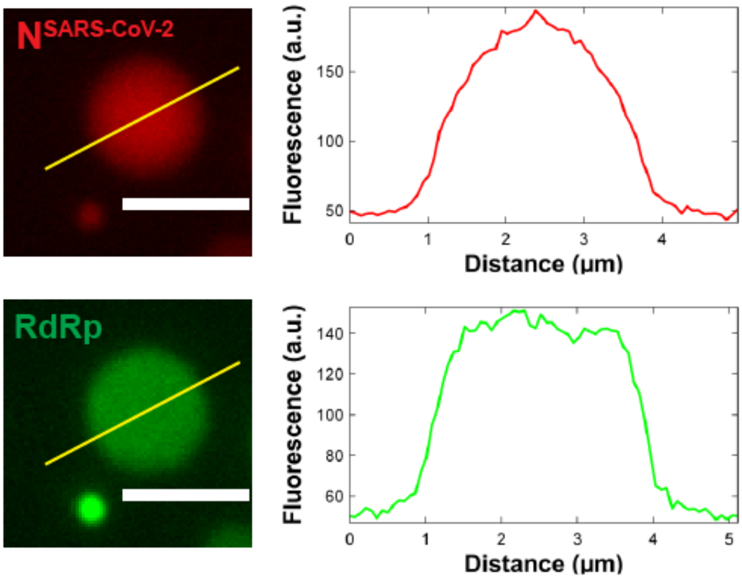


**Figure S6. Partitioning of the RdRp/RNA-complex into droplets formed by 50 µM N$^{SARS-CoV-2}$ and 1 µM polyU**

Fluorescence intensity measurement across the yellow line was plotted against the distance in micrometers. Strong enrichments of Alexa Fluor 594 fluorescently labeled N$^{SARS-CoV-2}$ and the RdRp-complex with a fluorescein-labeled minimal RNA hairpin template were observed inside the droplets in 20 mM NaPi, pH 7.5. Scale bar 3 µm.
